# Utanos: A general-purpose shallow whole-genome sequencing analysis workflow identifies interpretable copy number signatures

**DOI:** 10.64898/2026.07.22.739009

**Authors:** J Maxwell Douglas, Branden J Lynch, Jacky Chun Hei Yiu, Sheila Nicholson, Carlos Vasquez-Rios, Ding Ma, David G Huntsman, Yongjin P Park

**Affiliations:** Department of Basic and Translational Research, BC Cancer Research Institute, 675 West 10th Ave; Office 6-112, Vancouver, BC, V5Z 1L3; Department of Computer Science, ICICS/CS Building 201-2366 Main Mall, Vancouver, BC Canada V6T 1Z4; Department of Pathology and Laboratory Medicine, University of British Columbia, Vancouver, V5Z 1L3, Canada; Department of Gynecology and Obstetrics, University of British Columbia, Vancouver, BC, Canada; Department of Statistics, University of British Columbia, Vancouver, Canada

## Abstract

**Summary:** A modular FASTQ-to-figures solution for analyzing low-depth or ‘shallow’ whole-genome sequencing (sWGS) data. Shallow WGS can be used to detect copy number (CN) aberrations, Homologous Recombination Deficiency (HRD), and to detect and create CN signatures. Growing in popularity, this sequencing type is used for neonatal diagnostics and studying cancer. One of the major benefits is the reduced cost compared with deeper sequencing modalities such as Whole Genome Sequencing (WGS). With just 15 million reads often targeted, and sample substrate options like Formalin-Fixed, Paraffin-Embedded (FFPE) blocks widely available, this approach enables an affordable study to be performed at scale.

Our pipeline and R package are an end-to-end solution implemented with reusability and modularity in mind. It makes entry and exit from the ecosystem easy, providing regular standardized output formats throughout execution. The pipeline is written in the well-supported and cross-platform Nextflow framework and has been submitted for inclusion in nf-core. Additionally, a Docker image for the utanos R package has been created to improve modularity.

**Availability and Implementation:** The latest version of all software is freely available on GitHub. For the full processing pipeline, visit: https://github.com/Huntsmanlab/swgs-processing-pipeline. For just the utanos R package, visit: https://github.com/Huntsmanlab/utanos.

## 1 - INTRODUCTION

The era of high-throughput genomics has highlighted genomic aberrations in human pathologies like never before (Stratton *et al*. 2009, Chang *et al*. 2013). This is particularly consequential for cancer, a disease where genomic aberration is common (Stratton *et al*. 2009, Hanahan 2022). The gold standard of these genomic techniques for deep interrogation of the genome is whole-genome sequencing (WGS). It generates hundreds of gigabytes of output files and provides various types of subsequent analysis options (Stratton *et al*. 2009, Xiao *et al*. 2021, Bagger *et al*. 2024). However, WGS can quickly become less ideal when a targeted or shallower approach needs to be taken. Targeted panel sequencing (facilitated by amplicon-based methods) and, more recently, single-cell genomics (DLP+) are examples of technologies with clearly defined uses and advantages beyond WGS (Laks *et al*. 2019, Salehi *et al*. 2021). Single-cell genomics allows researchers to track cancer evolution at a clonal level (Salehi *et al*. 2021), and targeted panel sequencing enables SNP profiling in specific regions for a fraction of the cost of WGS (Tan *et al*. 2018).

Shallow WGS is another such technique. With low depth but broad coverage (Scheinin *et al*. 2014, Raman *et al*. 2019), it makes high-throughput whole-genome genomics more accessible. Unfortunately, because the field of genomic analysis did not evolve with the current model of sequencing as a sliding scale of depth, software tends not to cater well to this data type. Shallow WGS analysis is often tacked on to existing software as an extension, frequently WGS-focused tools, leaving the analyst to figure out which of the many parameters/modes/options are relevant vs. irrelevant.

We developed utanos and an accompanying shallow whole-genome sequencing pre-processing pipeline with this gap in mind. They were designed to make sWGS and related techniques more accessible to researchers and clinicians. With this toolset, a user can start from raw sequencing output and perform cohort-wide CN-profiling, create CN-signatures, call CN-signature exposures to existing signature sets, compare CN-Signatures, and detect homologous recombination deficiency (HRD). The utanos package is agnostic about the source of copy-number data, and so a user can begin using its features at any point of their analysis. A second benefit to being agnostic to CN-source is that utanos is also already well suited to analyze CNs coming from cutting-edge new technologies like cell-free DNA (cfDNA), and single-cell genomics such as DLP+ (Laks *et al*. 2019, Salehi *et al*. 2021). The use of utanos for DLP+ analysis is demonstrated in Supplemental Section 2.

## 2 - SOFTWARE OVERVIEW

The software described in this manuscript consists of two pieces. The first is a Nextflow pipeline (Di Tommaso *et al*. 2017) called shallowseq written in the style of ‘nf-core’ (Ewels *et al*. 2020), and the second is an R package called utanos. An exhaustive description of the modules/functions and their use is available on the respective websites for these two components. Here, we will describe just the core elements.

### 2.1 shallowseq

As seen in Figure 1A, the overall workflow can be divided into pre-processing and analysis.

**Figure 1.**
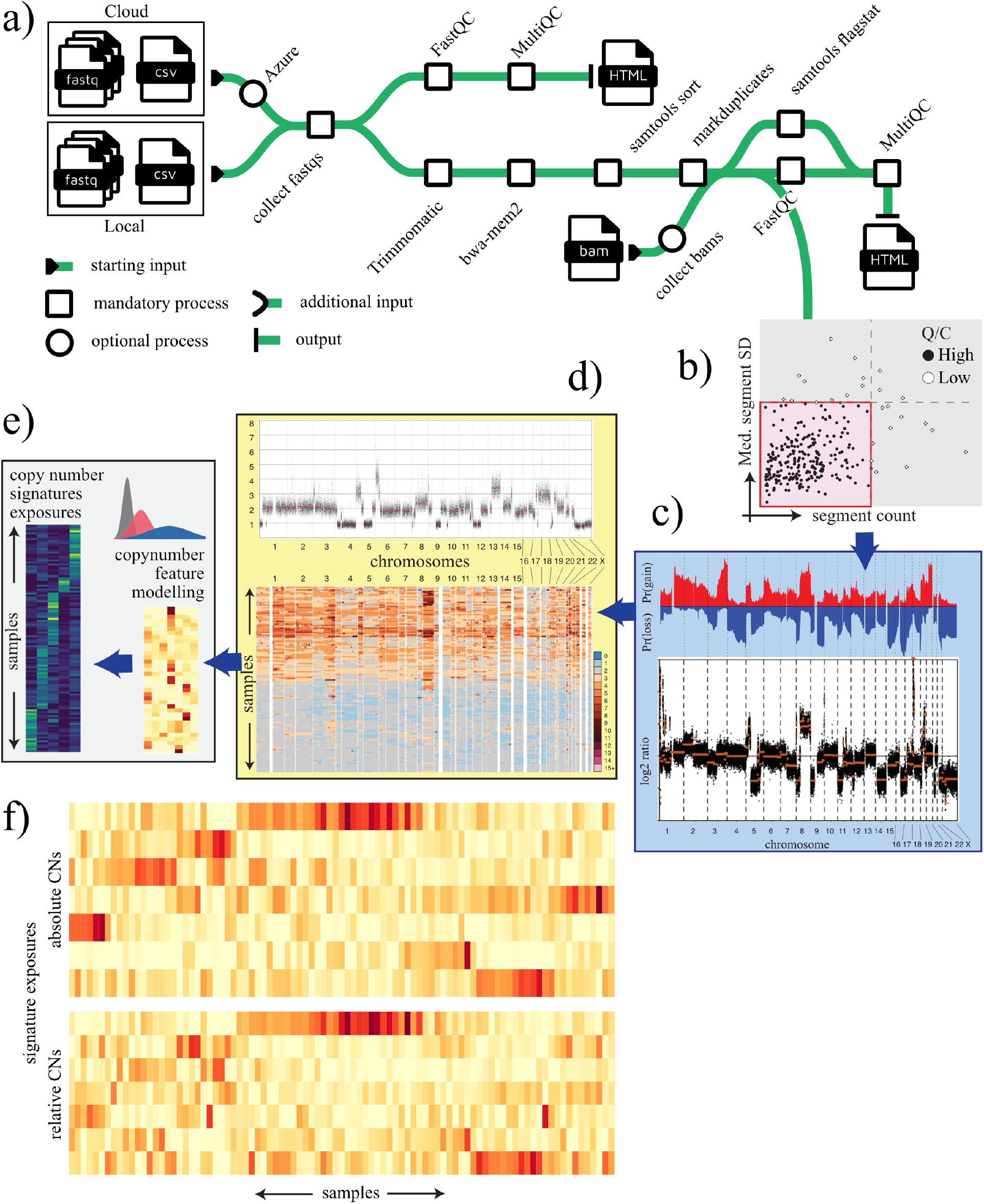
An overview of shallowseq and utanos. Utanos analysis begins at the relative copy-number calling stage with data filtration/quality-evaluation (b) and ends with copy-number signature comparisons (e). At most stages of an analysis workflow, utanos has functions available to visualize the data. These include heatmaps, copy-number scatterplots, and shaded line plots. The eight plots shown in b-e are not an exhaustive representation of the options. (f) Juxtaposed heatmaps of signature exposures by samples for both absolute (top) and relative (bottom) CN-Signatures.

#### Pre-processing

To begin pre-processing, Fastq/Bam files can be collected from the cloud or ingested from a local directory. The pipeline expects raw FASTQ or aligned BAM files from short-read sequencing (either single- or paired-end). A baseline quality assessment of the input reads is done using FastQC (Andrews 2010). Trimmomatic (Bolger *et al*. 2014) and bwa-mem2 (Vasimuddin *et al*. 2019) are used for trimming and alignment, respectively. Read sorting, indexing, and duplicate marking follow the alignment step using Samtools (Li *et al*. 2009) and Picard (Broad Institute 2019). Pre-processing concludes with calculating several QC metrics using FastQC and Samtools, and aggregating them using MultiQC (Ewels *et al*. 2016) for read coverage, mapping details, and duplicate marking.

#### Analysis

Pre-processed BAM files are fed into one or more options to calculate relative copy numbers: QDNAseq (Scheinin *et al*. 2014), WisecondorX (Raman *et al*. 2019), or ichorCNA (Adalsteinsson *et al*. 2017) using the utanos R package. Lastly, relative copy numbers are scaled back to absolute copy numbers again using utanos.

This approach has all the expected benefits of a modularized workflow design. Processing and analysis modules can be easily swapped out for newer or different procedures, and it is adaptive — compute demand scales cleanly across various scenarios, including both cluster-based high-performance computing environments and a single machine. Our new modules can be easily copied to other workflows without incurring heavy dependency.

### 2.2 utanos

#### Quality Evaluation

The quality of copy-number data varies significantly, and assessing quality in a large cohort can be labour-intensive. Utanos implements a bin-level filtering function, followed by the GetSampleQualityDecision function to support this task. By default, thresholds on the median segment variance and the total segment count are used to make the decision.

#### Scaling

Copy-number data, consisting of a numeric value per genomic bin, can be explored in the relative or absolute space. Utanos enables both. To scale copy-number data, utanos uses the method introduced in the ACE paper (Poell *et al*. 2019) and adapts code from the subsequent rascal package (Eldridge 2021). It calculates how well copy-number segments fit (the relative error) to integer copy numbers across a range of tumour purities/cellularities and ploidies and chooses the most likely solution (FindRascalSolutions and CalculateACNs functions).

#### CN-Summaries

Now that the CN data is streamlined, the bulk of the downstream analysis begins. Utanos enables the production of CN-summary plots, CN-summary tables, and CN-Diversity plots. The summary options display copy-number variation across a cohort of samples (Fig 1c), including an expanded version of a function found in the CGHbase package (van de Wiel and van Wieringen 2007). The CN-Diversity plot, rather than collapsing the data, plots each sample individually using a heatmap via the RCNDiversityPlot and ACNDiversityPlot functions (Fig. 1d). Both approaches offer filters to ease the iterative nature of investigating and discarding possible aberrations of interest (e.g., low-mappability or blacklisted regions, or commonly copy-number aberrant regions in the background human population (MacDonald *et al*. 2014)).

#### Feature Modelling and Creating Signatures

Modelling CN data represents the first step to creating and comparing CN signatures. To model copy-number aberrations, features (see Supplemental 1.1) within the CN-dataset are extracted using the ExtractCopyNumberFeatures function (code adapted directly from Macintyre *et al*. 2018) and modelled using flexmix (Grün and Leisch 2008) with mixtures of Gaussian or Poisson distributions. Sums-of-posteriors are calculated across features for each mixture using the flexmix package into a matrix; matrix decomposition is performed using the NMF package (Gaujoux and Seoighe 2010). The optimal rank is the number of discovered signatures. In utanos, the component-by-signature matrix is referred to as a ‘signature set’ or the ‘CN-signatures’.

#### Comparing Signatures

In addition to generating new CN-signatures, utanos includes functions to evaluate existing CN-signatures against a set of samples and to compare the derived signature sets. The YAPSA R package (Huebschmann *et al*. 2021) is used to evaluate a sample against an existing CN-signature set, while the custom-built SEAlluvialPlot and WassDistancePlot functions implement signature-set-to-signature-set comparisons (enables component-wise comparison).

#### Predicting HRD

Utanos contains a heavily refactored (improved modularity and cohort-wide prediction) implementation of version one of shallowHRD (Eeckhoutte *et al*. 2020) to identify samples that could have a non-functional homologous recombination pathway. This can be run on relative CN data, post low-mappability region filtering, using RunShallowHRD.

## 3 - CASE STUDY

Absolute CN-Signatures are a clinically relevant finding in cancer research, and their observation across cancer subtypes suggests common molecular mechanisms (Macintyre *et al*. 2018, Jamieson *et al*. 2024). A recent article highlighted that absolute copy-numbers themselves are not always necessary to create predictive CN-Signatures (Drews *et al*. 2022). This raises the question: what predictive signatures could be created using relative CNs instead? Since absolute CNs can be difficult to calculate in instances where a tumour is not very clonal, or cellularity is low, would relative CN-Signatures be useful, and are they also predictive of overall survival (OS)? We use the highly CN-aberrant cancer high-grade serous ovarian (HGSOC) as our model.

We obtained shallow WGS (Macintyre *et al*. 2018), deep WGS (PCAWG Consortium 2020), and exome (Zack *et al*. 2013) HGSOC data (see the methods of Macintyre *et al*. 2018 for details) and applied utanos to extract relative CN-features, model them, and create CN-Signatures. As a preliminary step, we ordered the samples and then compared the heatmaps of extracted signature exposures between relative and absolute values in the shallow WGS dataset. A visual inspection of the weighting of each sample to each signature revealed a high degree of similarity (Fig. 1f), particularly for signatures 1, 2, and 7. We then calculated the correlation of different biologic indicators to each relative CN-signature (Sup. Fig. 1B), closely mirroring what was done in the ACN case in Macintyre *et al*. 2018. This effectively reproduced most of the significant findings from the ACN case excluding those for signatures 3 and 5 (Sup Section 1.3, Sup. Fig. 1B and C), which were not as similar. The dissimilarity is also evident in the distribution of exposures across samples for signatures 3 and 5 in Fig. 1f). Additionally, we added the indicator of HRD status courtesy of utanos’ built-in tool. HRD status was found to be anti-correlated with signature 1 and positively correlated with signature 7 (Sup. Fig. 2). This is promising, given that signature 7 was identified as the non-BRCA HRD signature in the ACN case. Across the entire cohort, poor outcome was significantly predicted by CN signature 1 (p=0.0002), whilst good outcome was significantly predicted by exposure to CN signature 7 (p=0.008) (supplemental data section 3). We find, then, that RCN signatures can effectively recapitulate recreate most of the signatures derivable from ACNs.

Some recent research in a different modelling framework (Wang *et al*. 2021) has suggested that additional CN features beyond the original six are helpful. To test this for the relative CN case, we added two features as described in that paper: “nc50” (the minimum number of chromosomes needed to account for 50% of CN changes in a sample) and “cdist” (the scaled distance in base pairs of each breakpoint to the centromere, between 0 and 1). Additional feature modelling shows very modest improvements to the correlations between deep WGS/exome and shallow RCN-Signatures (Supplemental Figure 4C). The correlations between signatures and biologic indicators remain largely unchanged (Supplemental Figure 4B), and poor outcome was again significantly predicted by CN signature 1 (p=0.0001) whilst good outcome was significantly predicted by exposures to CN signature 7 (p=0.03) (Sup. Section 3).

## 4 - DISCUSSION

The utanos package and wrapping shallow WGS pipeline offer robust functionality for users interested in detecting and analyzing copy-number aberrations in genomic data. We adhere to good open-source practices, such as the FAIR principles for research software (Barker *et al*. 2022), and provide detailed vignettes in the linked repositories. Besides the case study just recounted, utanos has been used in several other projects.

1. The full pipeline and utanos package were used in a 2024 publication to discover copy-number signatures in p53-abnormal endometrial carcinoma (Jamieson *et al*. 2024).
2. Two small in-house projects involving single-cell DNA sequencing. A - to identify existing copy-number signatures created in a different modality. B - to identify novel cisplatin treatment signatures. See supplemental section 2 and supplemental figures 5-7.

A main limitation of our software is the scope. We designed the methods and structure specifically for shallow genomic sequencing, and therefore it focuses heavily on copy-number alterations. Increasingly, many types of research can rely on shallow sequencing; new statistical analyses will emerge, creating opportunities to expand the analytical component of our software, utanos, into other domains. For a more detailed exploration of the limitations and opportunities for expansion, see the supplementary material.

## AUTHOR CONTRIBUTIONS

J Maxwell Douglas: Conceptualization, Formal analysis, Methodology, Investigation, Software, Visualization, Validation, Writing —original draft. Branden J Lynch: Formal analysis, Methodology, Software, Visualization, Writing—original draft. Jacky Chun-Hei Yiu: Formal analysis, Visualization, Software, Writing—review & editing. Sheila Nicholson: Software, Validation. Carlos Vasquez-Rios: Software. Ding Ma: Software. David Huntsman: Funding Acquisition, Supervision. Yongjin-Park: Funding Acquisition, Supervision, writing – original draft.

## ACKNOWLEDGEMENTS

We’d also like to thank Sameer Shankar and Nirupama Tamvada for their efforts testing usage and building the utanos package.

