## Supplementary for "Utanos: A general-purpose shallow whole-genome sequencing analysis workflow identifies interpretable copy number signatures"

#### Table of Contents

**Section 1 – Main text supplemental ... 2**

Section 1.1 – Copy-number feature modelling ... 2

Section 1.2 – RCN-derived signatures effectively re-constitute the majority of ACN Signatures ... 3

Section 1.3 – Survival Analysis for HGSOC Relative CN-Signatures ... 6

Section 1.4 – Advantages and limitations of the shallowseq pipeline and utanos ... 7

**Section 2 – Secondary case study: detecting and creating CN-signatures in single-cell DNA sequencing ... 9**

Section 2.1 – Utanos can detect existing copy-number signatures in single-cell DNA sequencing libraries. 9

Section 2.2 – Utanos visualization functions can compare copy-number feature modelling of shallow vs. single-cell DNA sequencing to reveal overfitting ... 11

Section 2.3 – Utanos can create new copy-number signatures from TNBC single-cell DNA sequencing correlated with cisplatin treatment ... 13

### Section 1 – Main text supplemental

#### Section 1.1 – Copy-number feature modelling

Feature modelling was done with one of two feature sets: the standard set, comprising 6 copy-number features, or with 2 additional optional features.

Standard features:

| CN-Feature Name | Description |
|---|---|
| copynumber | Absolute copy-numbers at each genomic bin |
| changepoint | The magnitude (in CNs) of each copy number change |
| segsize | The genomic length (in bp) of each segment |
| brchrmarm | The number of breakpoints per chromosome arm |
| bp10MB | The number of breakpoints per 10 megabases |
| osCN | The genomic length (in bp) of oscillating copy number regions |

Utanos has also incorporated a few optional features introduced by Wang et. al. in 2021:

| CN-Feature Name | Description |
|---|---|
| nc50 | Minimum number of chromosomes (a count) needed to account for 50% of CN changes in a sample |
| cdist | Distance in base pairs of each breakpoint to the centromere |

Modelling relative CN data is done much the same as with absolute CNs, with a few modifications to three of the features detailed below and in supplemental figure 1.

| | |
|---|---|
| copynumber | Relative values < 0 are set to zero, then log transformed. |
| changepoint | Relative values < 0 are set to zero, then log transformed and the absolute value of the difference between adjacent segments is taken. |
| osCN | Relative values < 0 are set to zero, then log transformed rounded to the nearest 0.1. Oscillations are defined by a sliding window of 3 adjacent segments. When the outer two 2 segments have the same CN and the middle segment differs, a counter is incremented, giving the length of an oscillating region. |

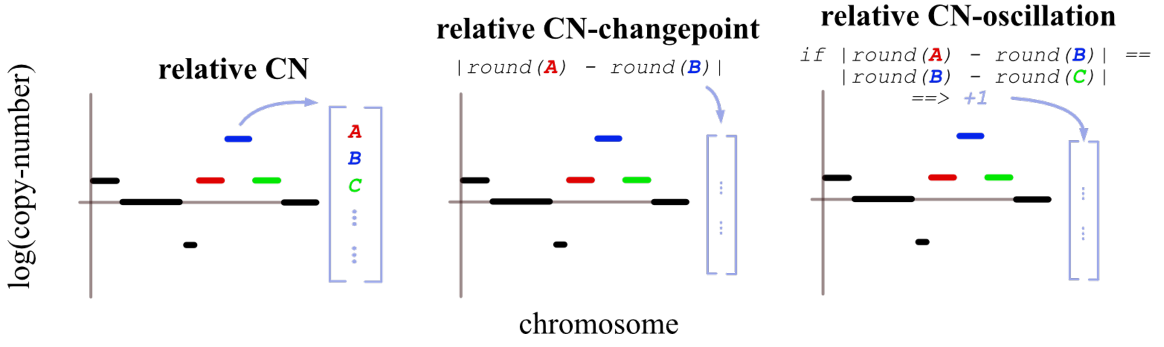


Supplemental Figure 1. Modelling CN, CN-changepoint, and CN-oscillation in relative CN data.

#### Section 1.2 – RCN-derived signatures effectively re-constitute the majority of ACN Signatures

After modelling and creating relative CN signatures for HGSOC, to evaluate the validity of these signatures we did a few tests and comparisons. As in Macintyre et. al. 2018 each relative CN signature, both using the standard feature set and with two additional features, was correlated with a range of clinical and molecular phenotypes (Sup. Fig. 2A). These metrics included the tandem duplicator score (TDP score), the telomere length, the age at diagnosis with HGSOC, the number of chromothriptic-like events, amplification-associated foldbacks inversions (Amp FBI), and the homologous recombination deficiency status (HRD status). For more information on the first five of these metrics please read that 2018 paper. The HRD status was evaluated using Utanos' implementation of shallowHRD, more details for which can be found on the package webpage.

a)

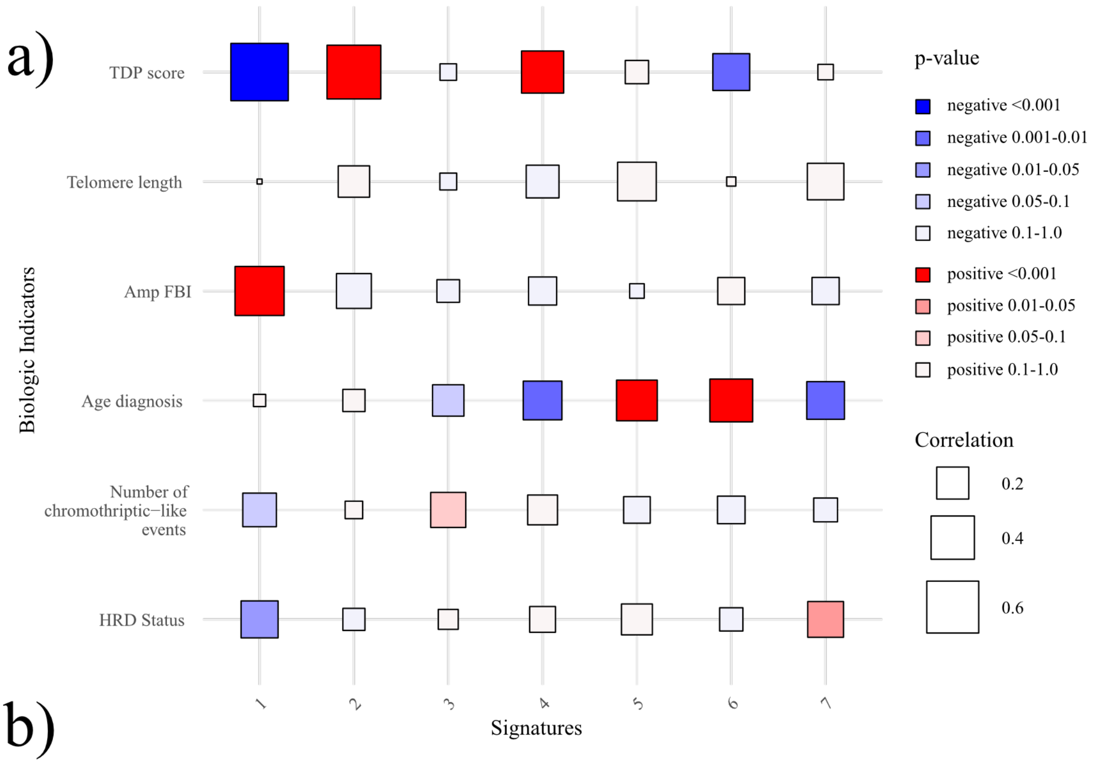


b)

| | 1 | 2 | 3 | 4 | 5 | 6 | 7 |
|:---|:---|:---|:---|:---|:---|:---|:---|
| PCAWG correlation | 0.91 | 0.80 | 0.07 | 0.74 | 0.51 | 0.75 | 0.77 |
| PCAWG p-value | 4e-12 | 2e-07 | 0.7 | 2e-06 | 0.003 | 1e-06 | 6e-07 |
| TCGA correlation | 1.00 | 0.88 | 0.55 | 0.85 | 0.30 | 0.81 | 0.91 |
| TCGA p-value | 1e-40 | 9e-11 | 0.001 | 1e-09 | 0.1 | 3e-08 | 1e-12 |

Supplemental Figure 2. A) Relative CN-Signature correlations with select molecular and clinical phenotypes. B) Correlations between relative CN-Signatures created using shallow WGS CNs and those created using data from two other sources: PCAWG and TCGA.

To validate the replicability of these relative CN-signatures we re-calculated the signatures in two other datasets using the models from shallow sequencing (shallow sequencing dataset: BritROC). These two different copy-number datasets came from whole genome sequencing (PCAWG Consortium 2020) and exome sequencing (Zack *et al.* 2013). Side-by side heatmaps of the loadings for each component composing the signature demonstrate that the first, second, fourth, sixth, and seventh signatures are quite similar (Sup. Fig. 3), and we confirmed this quantitatively (Sup. Fig. 2B). The eight feature signatures were subjected to the same test, and the results were much the same (Sup. Fig. 4).

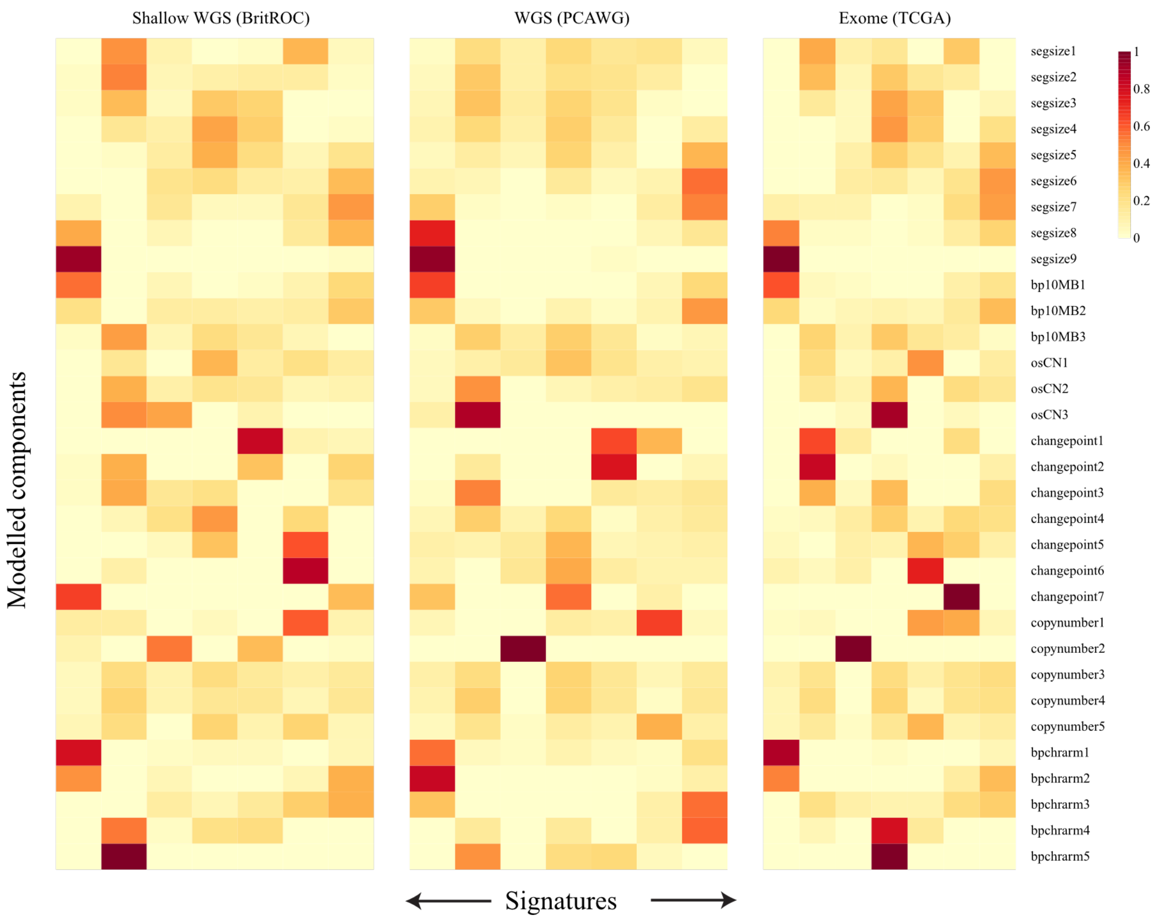


Supplemental Figure 3. Copy-number signatures calculated using three different datasets of high grade serous ovarian cancer samples. All three used copy-number feature models created from the shallow WGS sample set. On the vertical axis are the CN-feature model components, and on the x-axis are each signature.

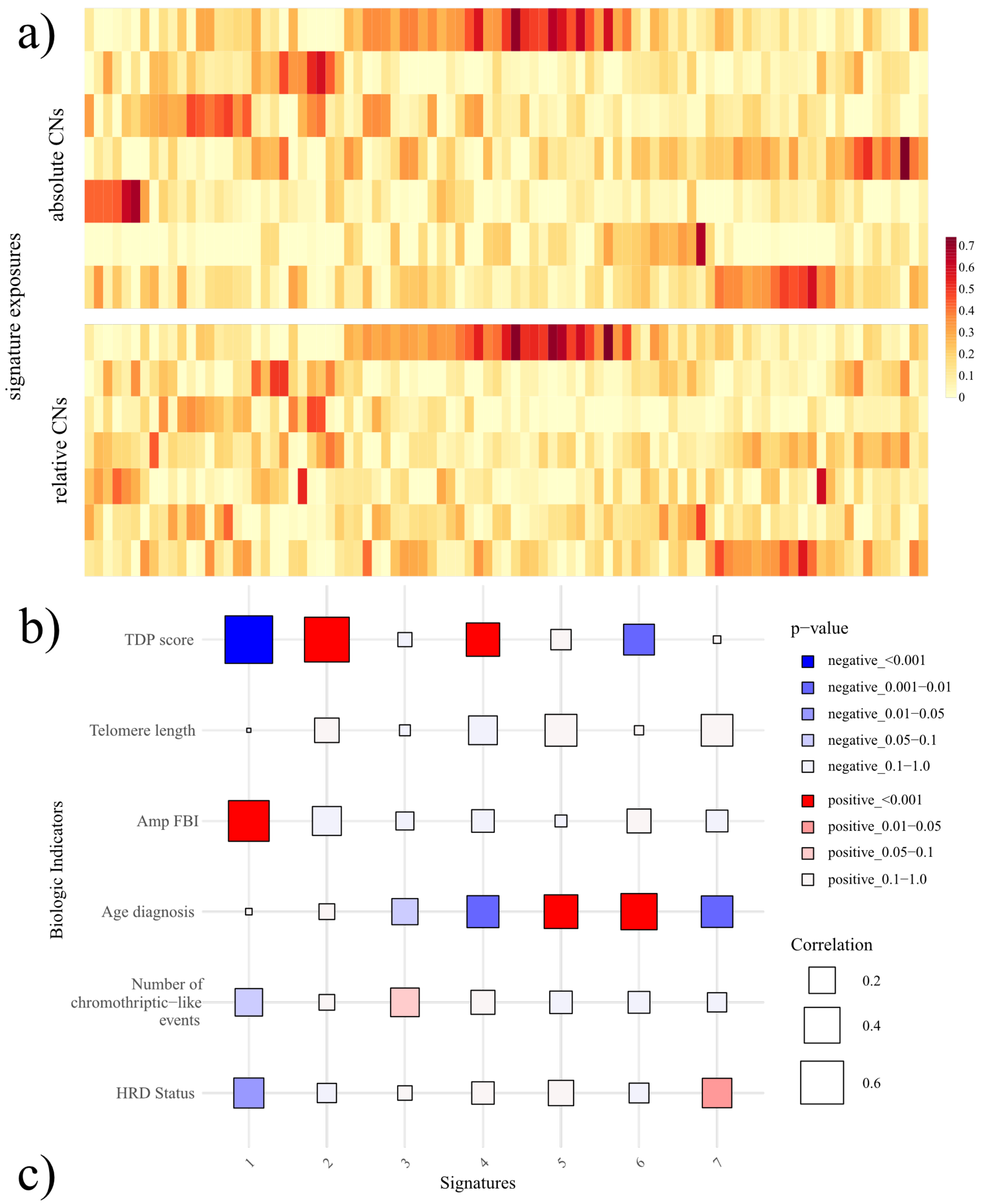


c)

| | 1 | 2 | 3 | 4 | 5 | 6 | 7 |
|---|---|---|---|---|---|---|---|
| **PCAWG correlation** | 0.90 | 0.86 | 0.25 | 0.78 | 0.66 | 0.73 | 0.79 |
| **PCAWG p-value** | 6e-13 | 1e-10 | 0.1 | 4e-08 | 1e-05 | 6e-07 | 2e-08 |
| **TCGA correlation** | 1.00 | 0.91 | 0.60 | 0.76 | 0.54 | 0.85 | 0.92 |
| **TCGA p-value** | 2e-45 | 2e-14 | 1e-04 | 1e-07 | 7e-04 | 7e-11 | 1e-14 |

Supplemental Figure 4. This set of panels is a mirror of Fig 1 f) and Sup. Fig. 2 just using the two additional CN-features. A) B) Relative CN-Signatures using 8 CN-features correlated with select molecular and clinical phenotypes. C) Correlations between relative CN-Signatures created using shallow WGS CNs and those created using data from two other sources: PCAWG and TCGA.

#### Section 1.3 – Survival Analysis for HGSOC Relative CN-Signatures

Using the survival package in R, cox-proportional hazards tests were performed for relative CN-signatures identified from shallow WGS HGSOC samples leaving out the signature with the lowest variance. Two sets of features were tested. In both cases (CN-Signatures modelled using six features (Code Output Box 1) or eight features (Code Output Box 2)) signatures 1 and 7 had $p < 0.05$. Signature 7 in both cases was predictive of better survival, whereas signature 1 was predictive of worse.

```
> summary(os_coxph)
Call:
coxph(formula = combined_os_survival ~ s1 + s2 + s4 + s5 + s6 +
    s7 + strata(study, age.cat, na.group = T), data = sig_data)

  n= 575, number of events= 335

         coef exp(coef)  se(coef)      z Pr(>|z|)
s1  0.138410  1.148446  0.037316  3.709 0.000208 ***
s2  0.060466  1.062332  0.055230  1.095 0.273606
s4 -0.001833  0.998169  0.046182 -0.040 0.968337
s5 -0.038427  0.962302  0.050806 -0.756 0.449440
s6  0.055290  1.056847  0.041489  1.333 0.182650
s7 -0.122981  0.884280  0.046383 -2.651 0.008015 **
---
Signif. codes:  0 '***' 0.001 '**' 0.01 '*' 0.05 '.' 0.1 ' ' 1

   exp(coef) exp(-coef) lower .95 upper .95
s1    1.1484     0.8707    1.0674    1.2356
s2    1.0623     0.9413    0.9533    1.1838
s4    0.9982     1.0018    0.9118    1.0927
s5    0.9623     1.0392    0.8711    1.0631
s6    1.0568     0.9462    0.9743    1.1464
s7    0.8843     1.1309    0.8074    0.9684

Concordance= 0.551  (se = 0.022 )
Likelihood ratio test= 23.38  on 6 df,   p=7e-04
Wald test            = 22.95  on 6 df,   p=8e-04
Score (logrank) test = 23.29  on 6 df,   p=7e-04
```

Code Box 1. Cox Proportional Hazards output for CN-Signatures modelled using six features.

```
 > summary(os_coxph)
Call:
coxph(formula = combined_os_survival ~ s1 + s2 + s4 + s5 + s6 +
    s7 + strata(study, age.cat, na.group = T), data = sig_data)

  n= 575, number of events= 335

       coef exp(coef) se(coef)      z Pr(>|z|)
s1  0.14424   1.15516  0.03767  3.829 0.000129 ***
s2  0.04015   1.04096  0.05511  0.728 0.466341
s4 -0.04858   0.95258  0.05002 -0.971 0.331451
s5  0.01828   1.01845  0.04865  0.376 0.707046
s6  0.03934   1.04012  0.04075  0.965 0.334376
s7 -0.10399   0.90123  0.04910 -2.118 0.034189 *
---
Signif. codes:  0 '***' 0.001 '**' 0.01 '*' 0.05 '.' 0.1 ' ' 1

   exp(coef) exp(-coef) lower .95 upper .95
s1    1.1552     0.8657    1.0729    1.2437
s2    1.0410     0.9606    0.9344    1.1597
s4    0.9526     1.0498    0.8636    1.0507
s5    1.0185     0.9819    0.9258    1.1204
s6    1.0401     0.9614    0.9603    1.1266
s7    0.9012     1.1096    0.8185    0.9923

Concordance= 0.545  (se = 0.023 )
Likelihood ratio test= 22.84  on 6 df,   p=9e-04
Wald test            = 22.37  on 6 df,   p=0.001
Score (logrank) test = 22.71  on 6 df,   p=9e-04
```

Code Box 2. Cox Proportional Hazards output for CN-Signatures modelled using six features.

#### Section 1.4 – Advantages and limitations of the shallowseq pipeline and utanos

Utanos' ability to allow the user to tune the width of CN bins is one of the key features that gives it broad use across sequencing modalities. Some perspective: newer methods such as DLP+ for single cell DNA sequencing most commonly have a bin width of 500kb (Laks et al. 2019) meanwhile CN-calling done on exome or whole-genome sequencing can be down to base-pair granularity. Shallow WGS derived from older FFPE blocks has an optimal window sizes that sits in the middle of these two modalites but can yield sequencing reads of highly varying quality. Quality variation in FFPE-derived sequencing means that for a given sequencing depth, bin widths of anywhere from 10kb to 500kb may be optimal. Software utilities that help users cleanly harmonize these differing dataset details is an understated but important element in making utanos useful.

Bin-size is a good place to begin detailing the limitations of our software. In genomic analysis bin-size is an artificial construct of convenience that depends on the quality, depth, and the platform of the experiment. As the granularity drops closer to bp-level resolution the applicability of our software diminishes and tools designed for WGS are better suited in order to take advantage of the additional information. In its current design, the pipeline + utanos are limited to the copy-number space, so a clear opportunity for extension would be shifting to include a broader set of structural variation (ex. inversions and transpositions).

While they were developed in-tandem, the target use-cases of the utanos package and processing pipeline do differ slightly. The whole ecosystem is dedicated to genomic copy-number, but CNs can be generated from several sequencing sources. So where the downstream analysis package utanos is sequencing-source agnostic, the processing pipeline that precedes it is limited to just shallow whole genome sequencing. An opportunity for extension is making the pipeline capable of processing other sequencing types for which copy-number the main data-type created (ex. Exome or single-cell DNA sequencing).

Utanos is able to do a rough job of detecting chromothripsis within the signature creation and detection modules via a combination of the oscillating copy-number and segment size CN-features. However, there isn't a standalone function that detects chromothripsis. This module would be akin to what we implemented for HRD detection by re-factoring shallowHRD (Eeckhoutte *et al*. 2020). Second, there is much potential in the space of comparing CN mutational signatures. In the last few decades there have been many methods published that attempt to create mutational signatures and papers that describe their use (Alexandrov *et al*. 2013, Funnell *et al*. 2019, Wang *et al*. 2021, Drews *et al.* 2022, Islam *et al.* 2022, and many more). However, given the complexity of modelling and the differences between one method and the next, knowing which may be more useful for a given dataset is challenging. Also challenging is explaining why, in a succinct way, one method behaves in a certain way compared to another. Utanos offers three or four functions/visualization options that allow users to compare the modelling of CN-features and signatures (MixtureModelPlots, WassDistancePlot, etc.).

To conclude, the pipeline and analysis package presented here offer a useful toolkit to the community for shallow genomic sequencing processing and copy-number analysis. The code is easily extensible building on previously published software in this space, it is modular, well documented, and offers flexible analysis options that can be mixed and matched. Accompanying these tools are visualization options that are publication ready or can be quickly modified to fit user needs.

### Section 2 – Secondary case study: detecting CN-signatures in single-cell DNA sequencing

#### Section 2.1 – Utanos can detect existing copy-number signatures in single-cell DNA sequencing libraries

To further demonstrate utanos' general applicability we ran three libraries of single cell DNA sequencing (DLP+) copy-numbers through signature exposure calling. These were HGSC patient-derived xenograft (PDX) samples published in 2022 by Funnell et al. In the 2022 paper, these PDX samples were categorised as belonging to the 'HRD-dup' strata using MMCTM (Funnell *et al*. 2019). This method ran on the corresponding whole genome sequencing samples and was based on both single-nucleotide and structural variants. In agreement with that previous result, as expected, the cells in these samples show higher exposures for the 2018 HGSOC CN-signatures: 1, 3, and 7. Signatures 3 and 7 are shown to be HRD-related in that paper, lending credence to the expectation that cancer signatures derived purely from copy-number data in one data modality capture a genomic profile common to other data modalities. This was shown in the 2018 paper for exome sequencing and WGS. We extend that here with single-cell DNA sequencing.

The extension is of technical interest given the different bin-sizes used in these two sequencing modalities. The 2018 CN-signatures created using shallow WGS used genomic bins of 30kb to call copy-numbers while the single-cell sequencing used bins of 500kb. Despite >10x difference in genomic resolution CN-signatures appear to identify common mutational profiles.

Utanos can also easily load, use, and evaluate CN-signatures created outside this ecosystem; all that's needed is a component-by-signature matrix. We loaded CN-signatures created from a pan-cancer cohort (Drews *et. al*. 2022), calculated exposures to the same scDNA HGSOC dataset, and plotted these in Supplementary Figure 5B. The cells also show higher exposures for signatures CX1 and CX3, while CX2 and CX5 are present to a lesser extent.

CX3 in the 2022 paper is observed to be associated with genetic perturbations beyond just the 'HRD' pathway, including genes like TP53. It makes sense for this signature to be elevated in our data as all three PDX samples also had TP53 mutations in the tumour from which they were derived. CX2 and CX5 were observed to be representative of a more moderate impairment of HR and thus their reduced prevalence in these cells is unsurprising.

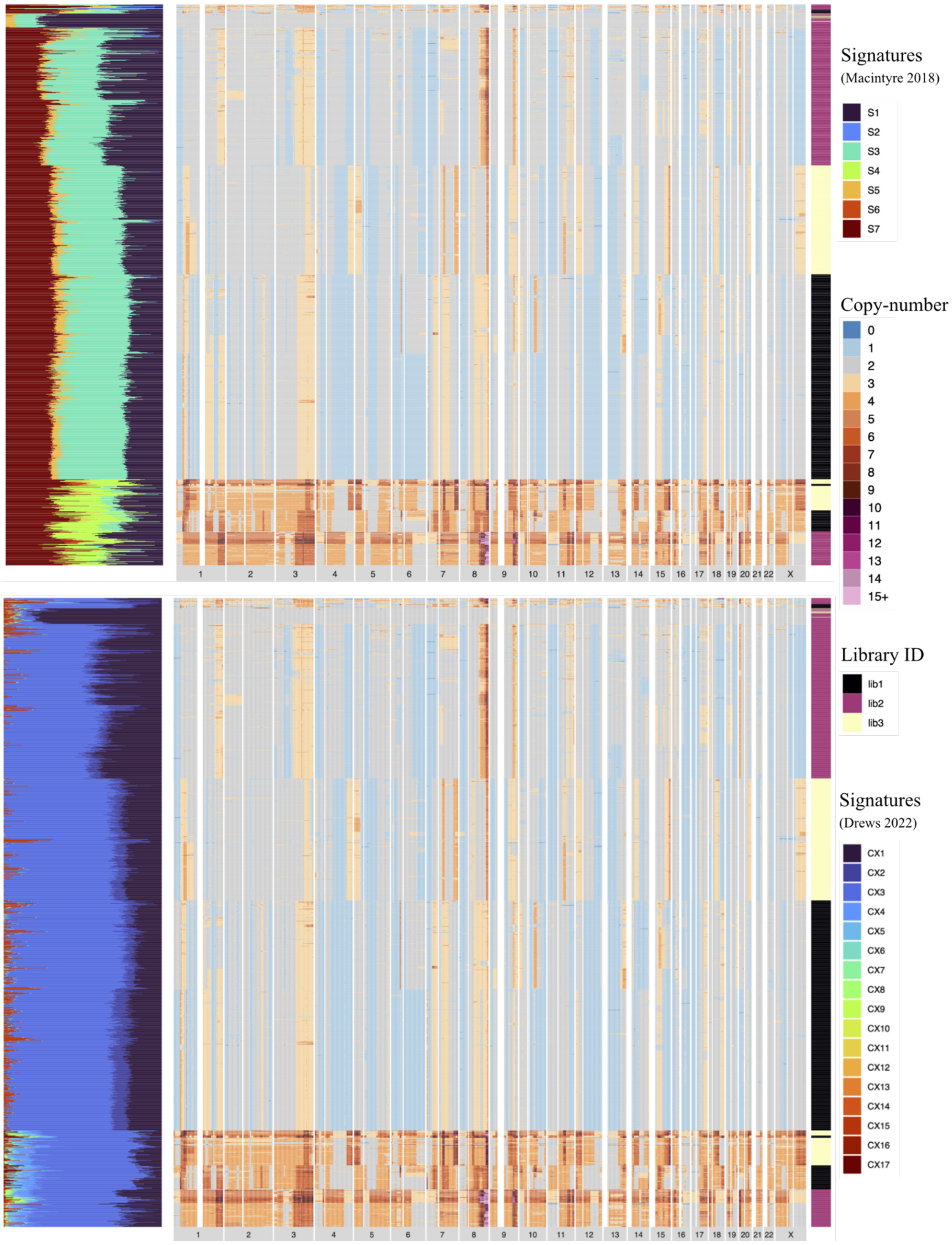


Supplemental Figure 5. CN-Diversity heatmaps of three combined DLP+ HGSC HRD samples (1356 cells total) juxtaposed with signature exposures. A) CN-Signatures created from shallow WGS HGSC samples. B) CN-Signatures created from the TCGA pan-cancer cohort and published in 2022 (Drews, 2022).

#### Section 2.2 – Utanos visualization functions can compare copy-number feature modelling of shallow vs. single-cell DNA sequencing to reveal overfitting

Modelling CN-features such as segment size (segsize) or the number of breakpoints per chromosome arm (bpchrarm) is done in utanos using mixtures of gaussian or poisson distributions. Each of these components can be compared to one another across datasets to examine the differences in structural variation. The WassDistancePlot function visualizes these mixture component differences as a heatmap meanwhile the GaussiansMixturePlot and PoissonsMixturePlot visualize them as line-plots.

To demonstrate the usage of these functions and explore the difference between scDNA and shallow WGS data from HGSOC, copy-number features were extracted and modelled in both modalities and are shown in Supplemental Figure 6. A parabolic arc is formed comparing segment sizes between data modalities (Sup. Fig. 6A), and an inverse of that comparing changepoints. The parabolic shape indicates that there are multiple scDNA segsize components that are more similar to individual shallow components than vice-versa for large segments while the reverse is true for smaller segment sizes.

This observation is repeated when comparing scDNA vs. pan-cancer exome CN-feature components and implies that there is less of a difference between the scDNA components vs. the shallow WGS or pan-cancer exome derived ones. Panels B and D for Supplemental Figure 6 specifically highlight how the CN mixture model components for single-cell tend to overlap more than the components from bulk data. Furthermore, the line-plot for changepoint from scDNA in Panel D contains two components at 1 that have nearly identical means and so differ by just their variance.

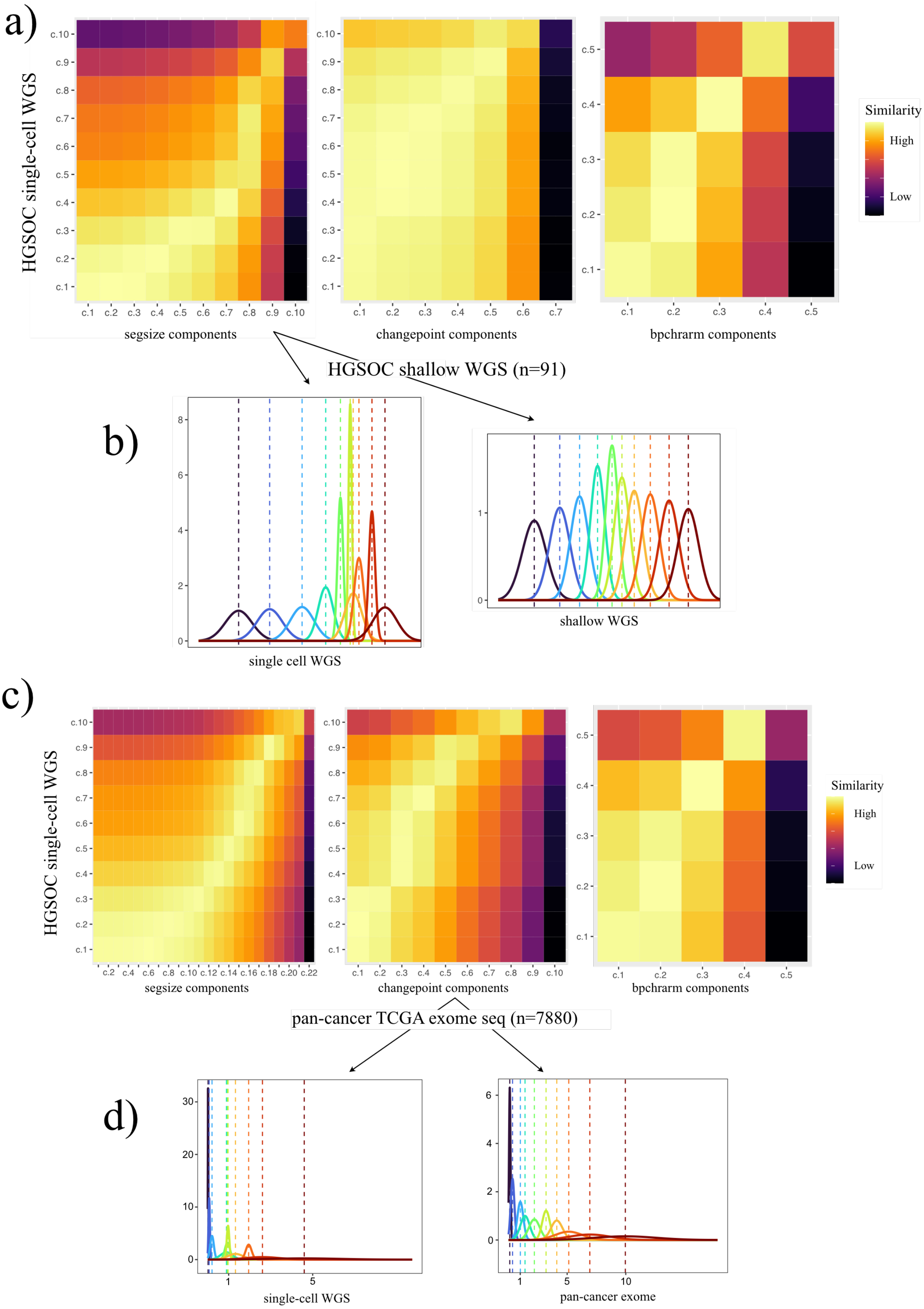


Supplemental Figure 6. Copy-number feature modelling of HGSOC using single-cell WGS and shallow WGS (panels A and B), and then single-cell WGS and TCGA pan-cancer exome sequencing (panels C and D). Panels A and C are heatmaps comparing CN-features modelled using a mixture of Gaussians. The similarity between components is determined using the Wasserstein Distance. Panels

B and D are line plots of the set of mixture components for a given CN-feature (segment size in B and changepoint in D) in a given dataset.

#### Section 2.3 – Utanos can identify new copy-number signatures from TNBC single-cell DNA sequencing correlated with cisplatin treatment

Utanos can also be used to create CN-Signatures in single-cell sequencing de-novo. To demonstrate, eleven triple-negative breast cancer (TNBC) DLP+ libraries were selected from Salehi et al. (2021) with the goal of identifying a signature for cisplatin treatment. We selected libraries based on the time point (passage/treatment cycle) and treatment status. Libraries that underwent treatment holidays were not included in the selection.

Two libraries were selected from passage 4 (Treatment Cycle X4); the study only introduced Cisplatin at X5, so both libraries are untreated. Two libraries were selected from passage 5 (Treatment Cycle X5); one library was treated with Cisplatin starting this passage. Four libraries were selected from passage 6 (Treatment Cycle X6); two were continuously treated with Cisplatin, and two remained untreated. Three libraries were selected from passage 8 (Treatment Cycle X8); two were continuously treated with Cisplatin, and one was untreated.

From the eleven libraries selected, we restricted to just four to create the signatures themselves (two untreated and two treated, with 2525 cells in total), and calculated signature exposures in the remaining seven. Utanos' signature creation procedure recognized six as the optimal number given cophenetic and dispersion metrics and a CN-Diversity plot is included as Sup. Fig. 7A visualizing these signature loadings adjacent to the cell CN data.

The calculated signatures showed significant association with treatment cycle (Sup. Fig. 7B). Across treatment cycles signature 1 decreased significantly while signatures 4 and 6 increased. Additionally, in treatment vs. no treatment scenarios, signatures 1 and 6 both showed significant differences again in opposing directions (Sup. Fig. 7C). Upon examination of signature 6, we noted a heavy loading on the third oscillating CN-feature component, pointing to a possible molecular effect of cisplatin.

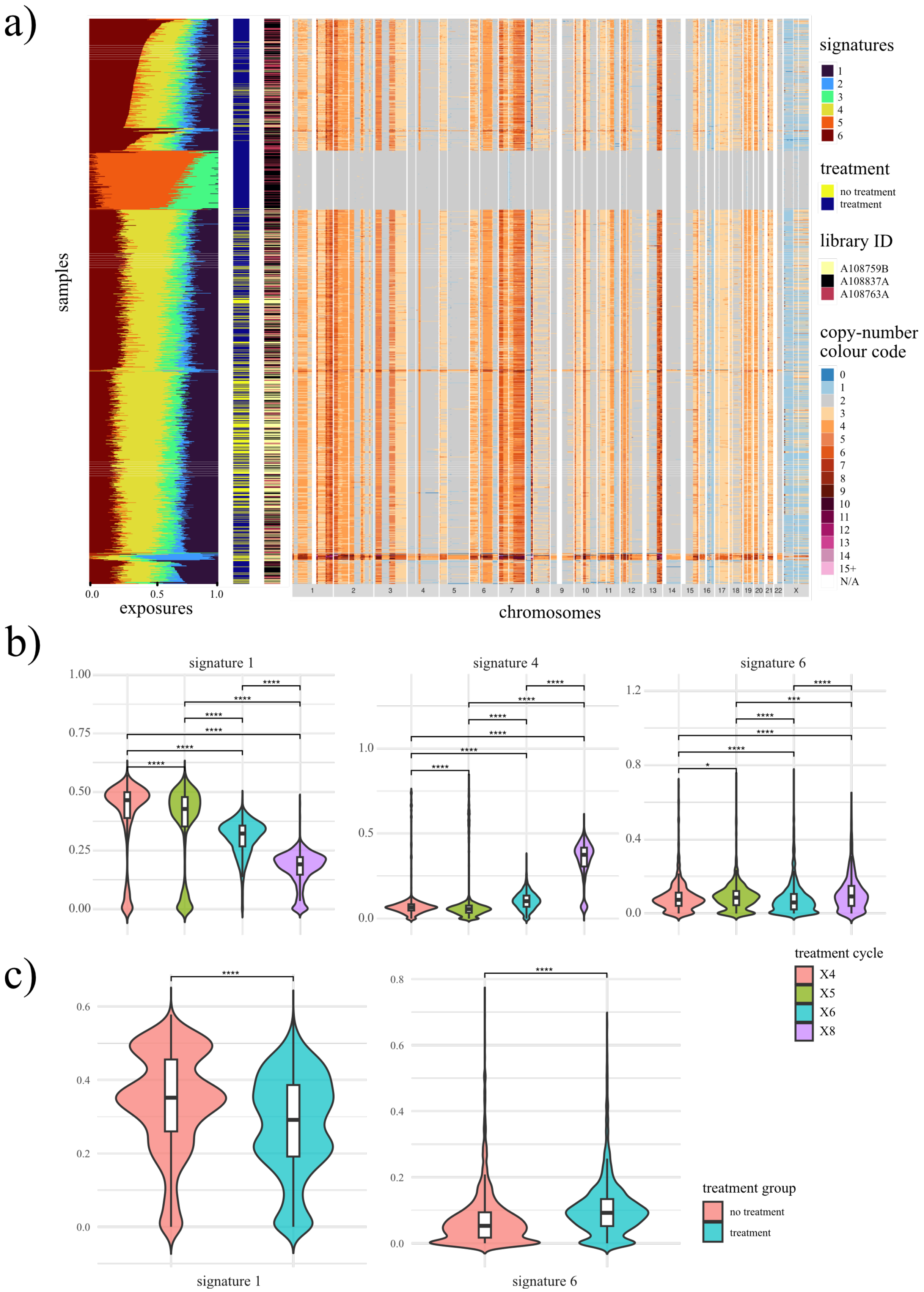


Supplemental Figure 7. CN-signatures of treated and untreated TNBC single cells. A) Juxtaposed CN-Diversity heatmap with annotation bars for the treatment status and library ID, and a stacked bar-plot of exposures for CN-signatures. Annotation bars and CN-diversity heatmap are row-ordered by the maximum signature exposure. B) Violin plots of signature exposure per-treatment cycle for three signatures. Pair-wise comparisons were done using a Wilcoxon test. C) Violin plots of signature exposure of treated vs. untreated cells. All cycles combined.
